# T cells expressing multiple co-inhibitory molecules in acute malaria are not exhausted but exert a suppressive function

**DOI:** 10.1101/2021.04.19.440453

**Authors:** Johannes Brandi, Cari Lehmann, Lea-Christina Kaminski, Julian Schulze zur Wiesch, Marylyn Addo, Michael Ramharter, Maria Mackroth, Thomas Jacobs, Mathias Riehn

## Abstract

Overwhelming activation of T cells in acute malaria is associated with severe outcomes. Thus, counter-regulation by anti-inflammatory mechanisms is indispensable for an optimal resolution of disease. Using *Plasmodium berghei* ANKA (PbA) infection of C57BL/6 mice, we performed a comprehensive analysis of co-inhibitory molecules expressed on CD4^+^ and CD8^+^ T cells using an unbiased cluster analysis approach. We identified similar T cell clusters co-expressing several co-inhibitory molecules like programmed cell death protein 1 (PD-1) and lymphocyte activation gene 3 (LAG-3) in the CD4^+^ and the CD8^+^ T cell compartment. Interestingly, despite expressing co-inhibitory molecules, which are associated with T cell exhaustion in chronic settings, these T cells were more functional compared to activated T cells that were negative for co-inhibitory molecules. However, T cells expressing high levels of PD-1 and LAG-3 also conferred suppressive capacity and thus resembled type I regulatory T cells. To our knowledge, this is the first description of malaria-induced CD8^+^ T cells with suppressive capacity. Importantly, we found an induction of T cells with a similar co-inhibitory rich phenotype in *Plasmodium falciparum* infected patients. In conclusion, we demonstrate that malaria-induced T cells expressing co-inhibitory molecules are not exhausted, but acquire additional suppressive capacity, which might represent an immune regulatory pathway to prevent further activation of T cells during acute malaria.

## Introduction

With 229 million cases and 409000 deaths worldwide in 2019, malaria remains a dangerous threat to humankind. 95 % of all malaria cases are due to an infection with *Plasmodium falciparum* (Global Malaria Programme: WHO Global, 2020). A liver stage and a subsequent blood stage characterize the infection.

There are two major characteristics of T cell responses to *Plasmodium* infections. On the one hand, there is severe immunopathology caused by T cells. Severe malaria cases are predominantly described during the first years of contact with the parasite and are reduced after repeated malaria episodes (Cowman et al., 2016). On the other hand, there is a lack of a protective memory response. Multiple malaria episodes do not lead to long-lasting protective immunity. Hepatocytes present pathogen-derived antigens during the liver stage of the infection, which opens a window of opportunity for adaptive immunity to interfere. However, an immune response against the liver stage during the first infections fails to deliver long-lasting protection. T cells, responsible for the local anti-malarial response, can be detected in the liver after vaccination in murine models, but do not persist for more than one month (Holz et al., 2018).

In mouse models of severe malaria, there is strong evidence that T cells are responsible for immunopathology (Howland et al., 2015). In humans, we and others could recently show that the severity of cerebral malaria correlates with the infiltration of CD8^+^ T cells to the brain (Riggle et al., 2020), and severe malaria is associated with an increase of granzyme B and activated cytotoxic T cells in the blood (Kaminski et al., 2019).

Interestingly, there is substantial evidence that T cell activation, induced by *Plasmodium* infection, concurs with the induction of co-inhibitory receptors (Biryukov & Stoute, 2014; Mackroth et al., 2016). Co-inhibitory or immune-modulatory receptors like PD-1, LAG-3, CD39, T cell immunoglobulin and mucin-domain containing-3 (TIM-3), and T cell immunoreceptor with Ig and ITIM domains (TIGIT) are primarily associated with a gradual dysfunction of T cells, called exhaustion (Blank et al., 2019). Exhaustion describes a mechanism to avoid tissue damage in organs under high antigen load (Cornberg et al., 2013).

Despite the high expression of co-inhibitory receptors in malaria (Mackroth et al., 2016), T cells mediate immunopathology and contribute to the development of malaria complications (Kaminski et al., 2019; Riggle et al., 2020), which is seemingly contradictory. Although a strong T cell response is initiated, a protective and long-lasting T cell memory is virtually absent in malaria.

We investigated these contradictions by avoiding the classic approach of analyzing malaria-specific T cell responses in a biased way. Using unbiased novel cluster analysis approaches, we identified the malaria-specific induction of distinct T cell populations with ambiguous properties. Apart from the expression of co-inhibitory receptors, such as PD-1, LAG-3, or TIM-3, the induced CD4^+^ and CD8^+^ T cells expressed high amounts of pro-inflammatory cytokines. For the first time, we also present that the same subsets of malaria-induced effector CD4^+^ and cytotoxic CD8^+^ T cells show suppressive capacity and can downregulate other CD4^+^ and CD8^+^ T cells. By identifying the dual function of malaria-specific T cells, we contribute to understanding the immunopathology in malaria.

## Results

### 1. Malaria-induced T cells express receptors associated with an exhaustion phenotype

Patients with acute malaria show different clinical pictures of malaria disease. Recently, we demonstrated that distinct expression patterns of T cell markers associated with immune regulation, such as CD39 and TIM-3, indicate less severe pathology during infection (Abel et al., 2018). Besides the activation of immunopathological T cells (Potter et al., 2006), we assume that the induction of immune-modulatory T cells occurs during infection with the parasite. To analyze the phenotype of malaria-induced T cells in the mouse model of PbA infection, we isolated T cells from different organs 6 days post infection (dpi). We analyzed the cells by employing an unbiased approach of self-organizing maps (SOM) followed by a minimum spanning tree (MST) and automated clustering to visualize and cluster possible T cell subsets (Van Gassen et al., 2015) (Fig. 1 & 2).

**FIG 1:**
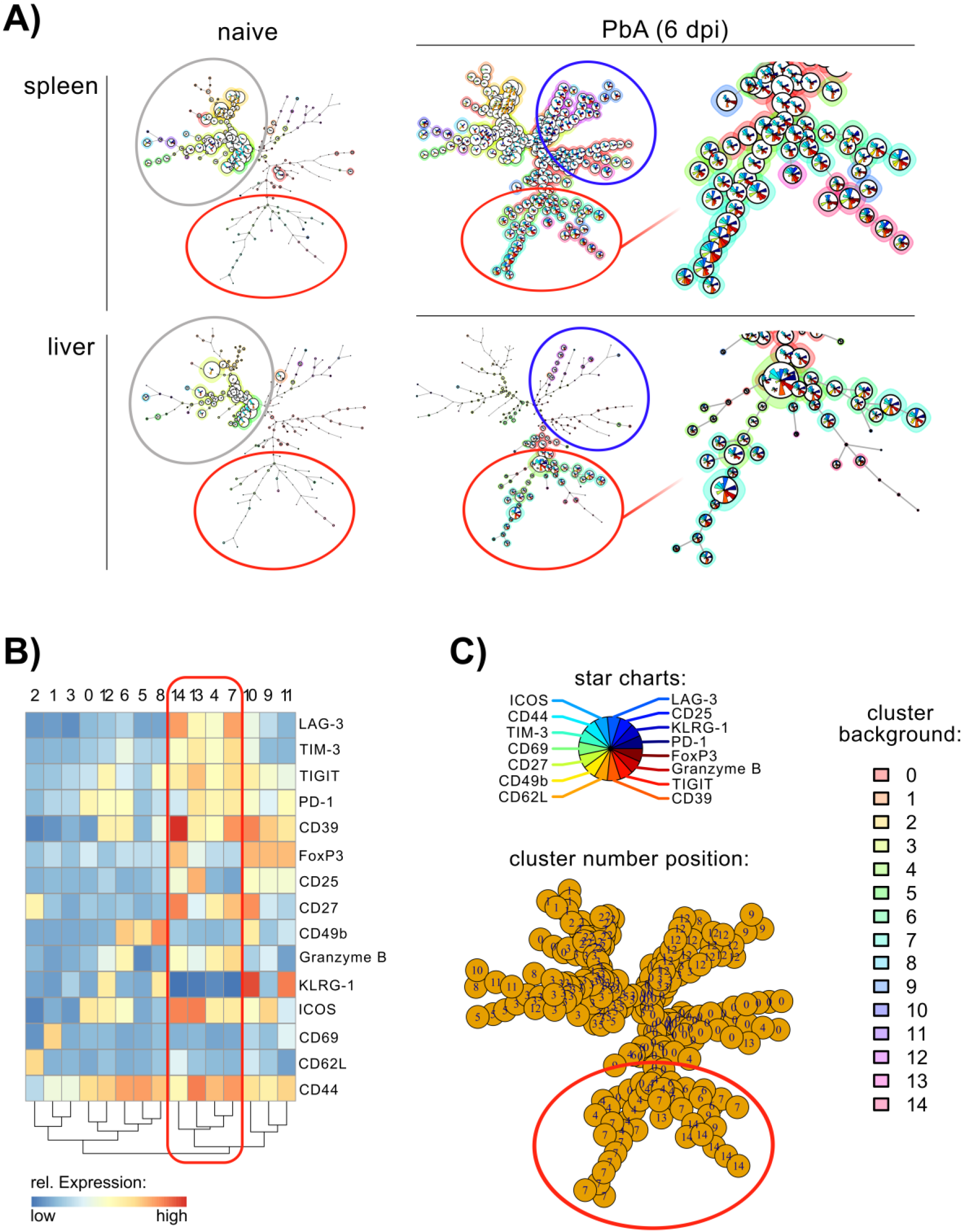
Malaria-induced CD4^+^ T cell subsets are distinguished by the expression of co-inhibitory molecules. **(A)** FlowSOM trees of CD4^+^ T cells from spleen and liver from naïve control and PbA-infected mice on day 6 post infection. Gray circle indicates T cells with naïve phenotype, blue circle indicates conventional activated T cells, red circle marks co-inhibitory-rich T cells. **(B)** Expression heatmap of indicated markers of each metacluster. **(C)** Legends for FlowSOM tree; star charts show color code for the expression of each marker per cluster; cluster number position depicts metacluster number and position in the tree, cluster background shows the background color of each metacluster in the FlowSOM tree.

**FIG 2:**
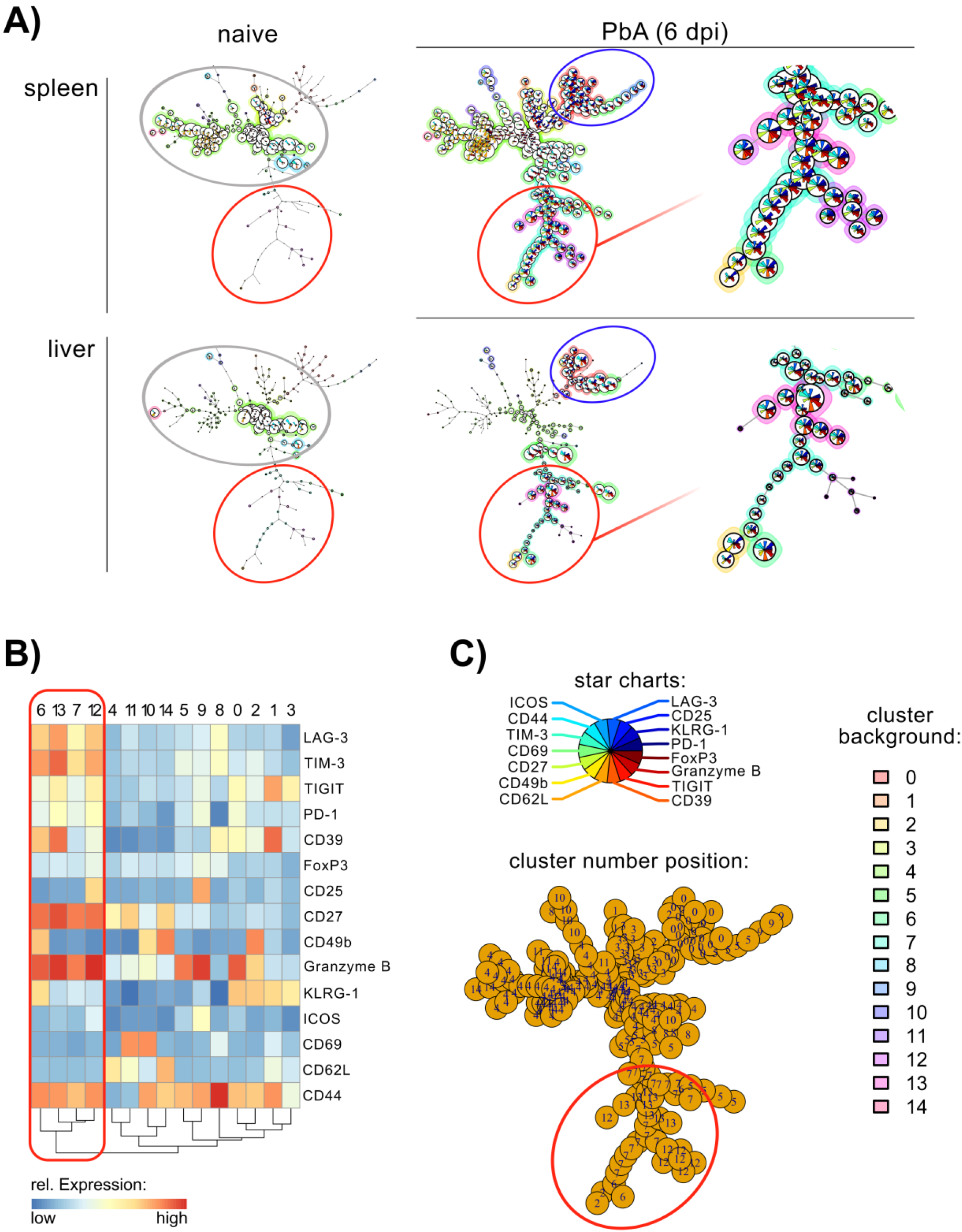
Malaria-induced CD8^+^ T cell subsets are distinguished by the expression of co-inhibitory molecules. **(A)** FlowSOM trees of CD8^+^CD3^+^ T cells from spleen and liver from naïve control or PbA-infected mice on 6 dpi. Gray circle indicates T cells with naïve phenotype, blue circle indicates conventional activated T cells, red circle marks co-inhibitory-rich T cells. **(B)** Expression heatmap of indicated markers of each metacluster. **(C)** Legends for FlowSOM tree; star charts show color code for the expression of each marker per cluster; cluster number position depicts metacluster number and position in the tree, cluster background shows the background color of each metacluster in the FlowSOM tree.

FlowSOM uses two-level clustering with a Minimum Spanning Tree as visualization (Van Gassen et al., 2015). The first level is based on an artificial neural network, the self-organizing map (SOM). Based on intracellular and surface protein expression (as indicated in Fig 1B and 2B), the SOM is trained to assemble similar cells into a cluster grid. The topological information of the trained grid is visualized in a minimum spanning tree (Fig 1A and 2A). The second level of clustering is indicated by the background color of the branches of the MST (Fig 1A & Fig 2A) as well as the cluster number (Fig 1C & Fig 2C). It is based on consensus hierarchical clustering as described previously (Van Gassen et al., 2015; Wilkerson & Hayes, 2010).

The artificial neural network is trained using isolated T cells from spleens of PbA-infected animals and naïve controls. We assume that T cells found in spleens of infected animals reflect most naïve and activated T cell subsets throughout the organism. Therefore, T cells from livers of animals isolated from the indicated groups are also analyzed with the trained algorithm. We analyzed CD4^+^ T cells and CD8^+^ T cells separately.

Using the computed scaffold for CD4^+^ T cells from the spleens of naïve animals, we identified distinct clusters of naïve T cells (Fig 1A, gray circle). These clusters belong mainly to metacluster 1, 2, and 3. Checking the indicated metacluster in the heatmap (Fig 1B) reveals that these cells lack the expression of receptors associated with activation, like CD44 or Inducible T cell Costimulator (ICOS). In addition, cluster 2 expresses high amounts of CD62L, which is a marker for naïve lymphocytes. We found a similar cluster and distribution of naïve T cells in the liver of uninfected mice (Fig 1A, gray circle), indicating the distribution of naïve T cells between lymphoid organs and organs in the periphery.

Using the scaffold on CD4^+^ T cells isolated from the liver of PbA infected animals, the induction of unique metaclusters can be seen. These induced T cell clusters can also be found in the spleens of infected animals but not in the liver or the spleens from naïve animals. Apart from a branch of classically activated CD44^+^CD62L^−^ T cells, we identified another branch of induced T cells (Fig 1A, red circle) which mainly consists of the metaclusters 14, 13, 4, and 7. CD4^+^ T cells from the metacluster 14 are CD25^+^FoxP3^+^ (Fig 1B) and can be considered classical regulatory T cells (Miyara & Sakaguchi, 2011). Metacluster 14 can be found in the spleen, but not the liver of infected mice. In the liver, metaclusters 7 and 4 are most abundant (Fig 1A). T cells represented in this cluster are FoxP3^−^ (Fig 1B). Apart from the high expression of CD44 or ICOS, which are associated with T cell activation (Hutloff et al., 1999), these clusters are characterized by immune regulatory molecules like TIM-3, CD39, TIGIT, PD-1, and LAG-3 (Fig 1B, red frame). Low amounts of the marker for finally differentiated T cells, Killer cell lectin-like receptor subfamily G member 1 (KLRG-1) (Sarkar et al., 2008), are detected. However, as granzyme B is expressed in these T cell subsets, we consider these T cells to be highly differentiated effector T cells.

Interestingly, by simultaneously analyzing CD8^+^ T cells, we recognized a similar T cell activation and differentiation pattern. Naïve CD8^+^ T cells were found in the spleen and liver of naïve mice. Frequencies of naïve cells are reduced after infection with PbA (Data not shown). Compared to naïve animals, the branches of malaria-induced CD8^+^ T cells dominate (Fig 2A, blue and red circle). Again, distinct clusters show the expression of co-inhibitory molecules like TIGIT, TIM-3, PD-1, and LAG-3 (Fig 2A, red circle, mainly Clusters 6, 7, 12 and 13). Furthermore, they are characterized by the expression of CD39 and granzyme B and the downregulation of KLRG1 (Fig 2B, red frame).

In conclusion, the data demonstrate that CD4^+^ and CD8^+^ T cells induced by PbA infection can be divided into two major groups. One group is positive for CD44 and KLRG1 but expresses low levels of co-inhibitory molecules. The second group expresses CD44 and LAG-3 but almost no KLRG1. More specifically, the latter subset is characterized by expression of a high amount of anti-inflammatory receptors, including co-inhibitory molecules (PD-1, LAG-3) or functional receptors associated with T cell exhaustion in chronic infection models (CD39)(Canale et al., 2018; Wherry, 2011). Strikingly, they also express the cytotoxic effector molecule granzyme B. Recently, LAG-3 was described as a reliable marker for induced potential regulatory B cells (Lino et al., 2018). Moreover, when co-expressed with CD49b, it is described as a marker for regulatory type 1 T cells (Gagliani et al., 2013).

### 2. Blood stage-induced LAG-3^+^ T cells are not exhausted

We observed the induction of T cells expressing high levels of LAG-3 and other co-inhibitory molecules by the PbA infection. To dissect immune responses to the liver and blood stage of infection, we experimented with 3 distinct treatment groups (Fig 3A). C57BL/6 mice in group 1 were infected with viable sporozoites. These mice undergo a complete liver and blood stage. Group 2 was infected with viable sporozoites and treated with anti-parasitic pyrimethamine on day 5 post infection. Pyrimethamine is toxic for extracellular parasites in the blood and shortens the blood stage (McGregor & Smith, 1952). Group 3 was infected with irradiated sporozoites. Irradiated sporozoites are able to infect hepatocytes, but they do not develop to schizonts and remain in the liver without provoking a blood stage (Silvie et al., 2002). Comparing the immune response to the liver and the blood stage, our data reveal that LAG-3 on CD8^+^ and CD4^+^ T cells was induced during the blood stage but not the liver stage of the disease (Fig 3C and Fig 4A). LAG-3^+^ T cells could be found in blood, spleen and in peripheral organs like the brain and liver (Fig 3C), indicating a ubiquitous distribution during the infection with PbA. Remarkably, when administering an anti-malarial treatment with pyrimethamine, which leads to delayed antigen withdrawal (experiment described in Fig 3A), LAG-3^+^ T cells disappeared rapidly (Fig 3C and 4A), indicating a transient, antigen-specific induction of LAG-3^+^ T cells.

**FIG 3:**
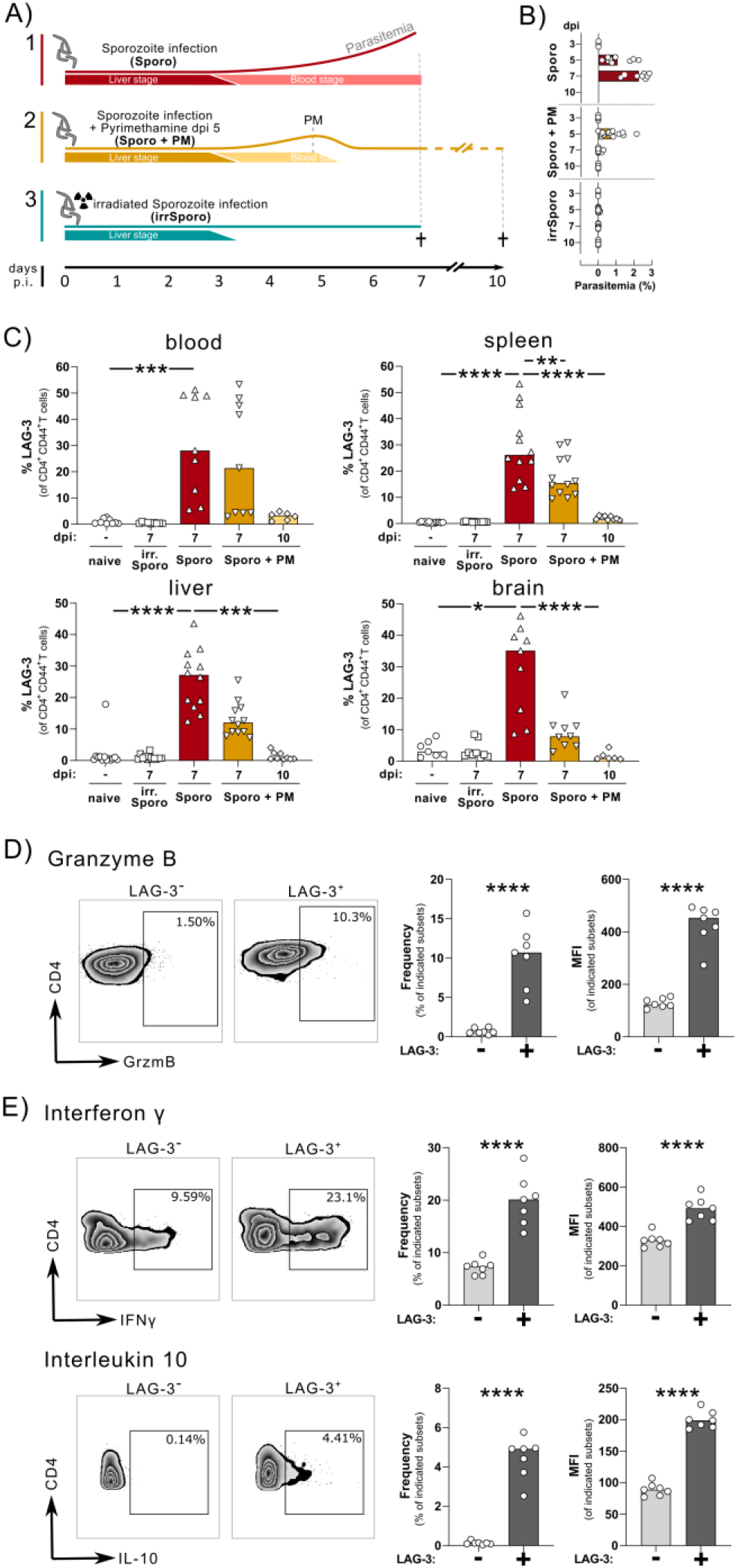
Blood stage-induced CD4^+^LAG-3^+^ T cells are not exhausted. **(A)** Experimental scheme to dissect impact on induced T cells of liver stage or blood stage of PbA infection. Group infected with viable sporozoites (Sporo) reflects liver and full blood stage (red). Group infected with viable sporozoites and treated with pyrimethamine (Sporo + PM) reflects a reduced blood stage (yellow). Group infected with irradiated sporozoites (irrSporo) are limited to a liver stage and do not show a blood stage (blue). **(B)** Blood parasitemia on indicated days post infection. **(C)** Frequency of LAG-3 expression by CD4^+^CD44^+^ T cells in indicated organs and groups. Kruskal-Wallis test with Dunn’s multiple comparisons test was used. For better readability, not all significances are depicted. **(D)** Frequency and Median Fluorescence Intensity (MFI) of granzyme B expression on LAG-3^+^ or LAG-3^−^ CD4^+^CD44^+^ T cells measured *ex vivo* 7 dpi. **(E)** Frequency and MFI of IL-10 and IFN_*γ*_ stained after 5 h Phorbol 12-Myristat 13-Acetat (PMA)/ Ionomycin (Iono) restimulation of T cells isolated 7 dpi. **(D)** and **(E)** Mann-Whitney test was applied. **(A-E)** P values ≤ 0.05 (*),

**FIG 4:**
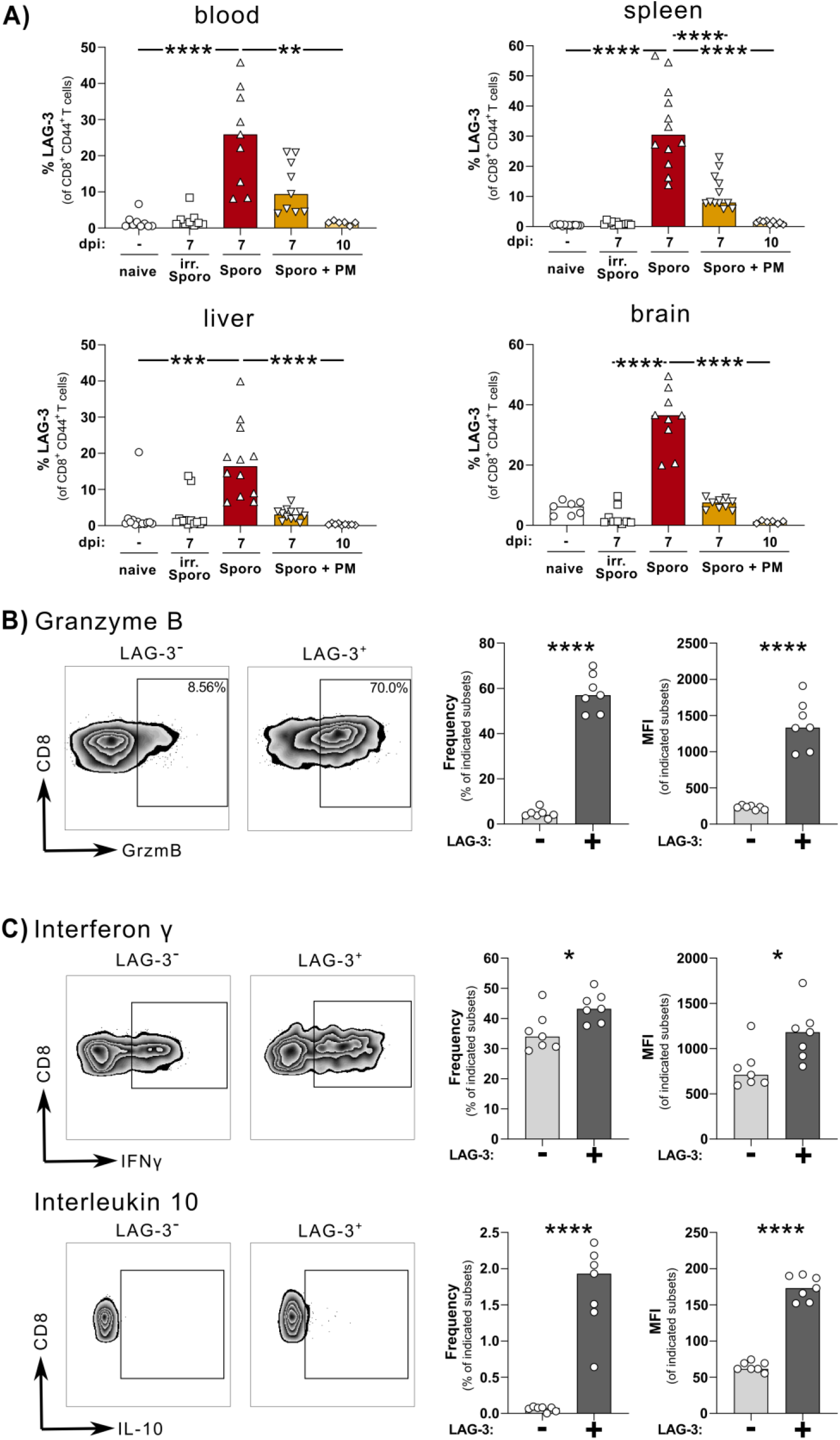
Blood stage-induced CD8^+^LAG-3^+^ T cells are not exhausted. **(A)** Frequency of LAG-3 expression by CD8^+^CD44^+^ T cells in indicated organs and groups. Kruskal-Wallis test with Dunn’s multiple comparisons test was used. For better readability, not all significances are depicted. **(B)** Granzyme B expression on LAG-3^+^ or LAG-3^−^ CD4^+^CD44^+^ T cells measured *ex vivo* on 7 dpi. **(C)** IL-10 and IFN_*γ*_ stained after 5 h PMA/Iono restimulation of T cells isolated on 7 dpi. **(B)** and **(C)** Mann-Whitney test was used. **(A-C)** P values ≤ 0.05 (*), ≤ 0.01 (**), ≤ 0.001 (***) or ≤ 0.0001 (****) were considered statistically significant.

We stained *ex vivo* for granzyme B in isolated T cells to validate that malaria-induced T cells are functional. We also detected the effector cytokine Interferon *γ* (IFN_*γ*_) and the anti-inflammatory cytokine IL-10 in restimulated splenocytes from infected animals. We observed that CD4^+^CD44^+^LAG-3^+^ T cells expressed more granzyme B and higher amounts of IFN_*γ*_ and IL-10 compared to activated CD4^+^CD44^+^LAG-3^−^ T cells (Fig 3D and E). Indeed, our results show that the expression of granzyme B and IL-10 by CD4^+^ T cells is associated with the expression of LAG-3 (Fig 3). This indicates that the expression of co-inhibitory receptors during acute malaria is not associated with exhaustion.

Furthermore, the frequency of CD8^+^CD44^+^LAG-3^+^ T cells was increased during the blood stage of infection. CD8^+^CD44^+^LAG-3^+^ T cells could be found in peripheral organs (liver, brain), secondary lymphatic tissue (spleen), and the blood (Fig 4A). *Ex vivo* staining of granzyme B revealed that CD8^+^CD44^+^LAG-3^+^ T cells produce more cytotoxic granzyme B than T cells not expressing LAG-3 (Fig 4B). Additionally, the expression of the effector cytokine IFN_*γ*_ and the anti-inflammatory cytokine IL-10 was increased in the subset of CD8^+^CD44^+^LAG-3^+^ T cells compared to their LAG-3^−^ counterparts (Fig 4C). Withdrawal of antigen by treatment with pyrimethamine led to a fast reduction of LAG-3^+^ T cells. On day 10 after infection or day five after pyrimethamine treatment, LAG-3 expression was sharply reduced and hard to detect in any organ (Fig 4A).

### 3. CD8^+^LAG-3^+^ and CD4^+^LAG-3^+^ T cells exert a suppressive capacity

Despite the expression of LAG-3 and other co-inhibitory molecules by malaria-induced T cells, our data imply that these cells are not exhausted but develop a dual phenotype with pro- and anti-inflammatory characteristics. Indeed, inducible Type 1 regulatory T cells (Tr1) are also described to be induced during malaria (Brockmann et al., 2018). Based on these results, showing suppressive capacity of CD4^+^LAG-3^+^ T cells, we hypothesize that malaria-induced CD8^+^LAG-3^+^ T cells are suppressive as well. To this end, we conducted *in vitro* suppression assays through co-culture of naïve CD4^+^ and CD8^+^ T cells with different subsets of malaria-induced T cells. Immuno-suppressive Tr1 cells were described as being CD4^+^CD49b^+^LAG-3^+^ T cells (Gagliani et al., 2013). Thus, we sorted malaria-induced CD4^+^CD49b^+^LAG-3^+^ T cells (described as Tr1 cells) and CD4^+^CD49b^−^LAG-3^+^ T cells to compare their suppressive capacity to CD49b^+^LAG-3^−^ T cells. Additionally, we performed suppression assays with the same LAG-3^+^ and/or CD49b^+^ subsets of CD8^+^ T cells.

First, we investigated whether malaria-induced CD4^+^ and CD8^+^ T cells can suppress the proliferation of naïve CD4^+^ T cells. CD4^+^LAG-3^+^ T cells suppressed the proliferation of CD4^+^ T cells in a dose-dependent manner (Fig 5A and B). We could not detect a difference in the regulatory capacity between LAG-3^+^CD49b^+^ T cells and LAG-3^+^CD49b^−^ T cells; both subsets suppressed proliferating CD4^+^ T cells equally. Like CD4^+^LAG-3^+^ T cells, malaria-induced CD8^+^LAG-3^+^ T cells could also suppress CD4^+^ T cells in a dose-dependent manner (Fig 5A and B). Again, the co-expression of CD49b by CD8^+^LAG-3^+^ T cells was not associated with a higher suppressive capacity (Fig 5A and B).

**FIG 5:**
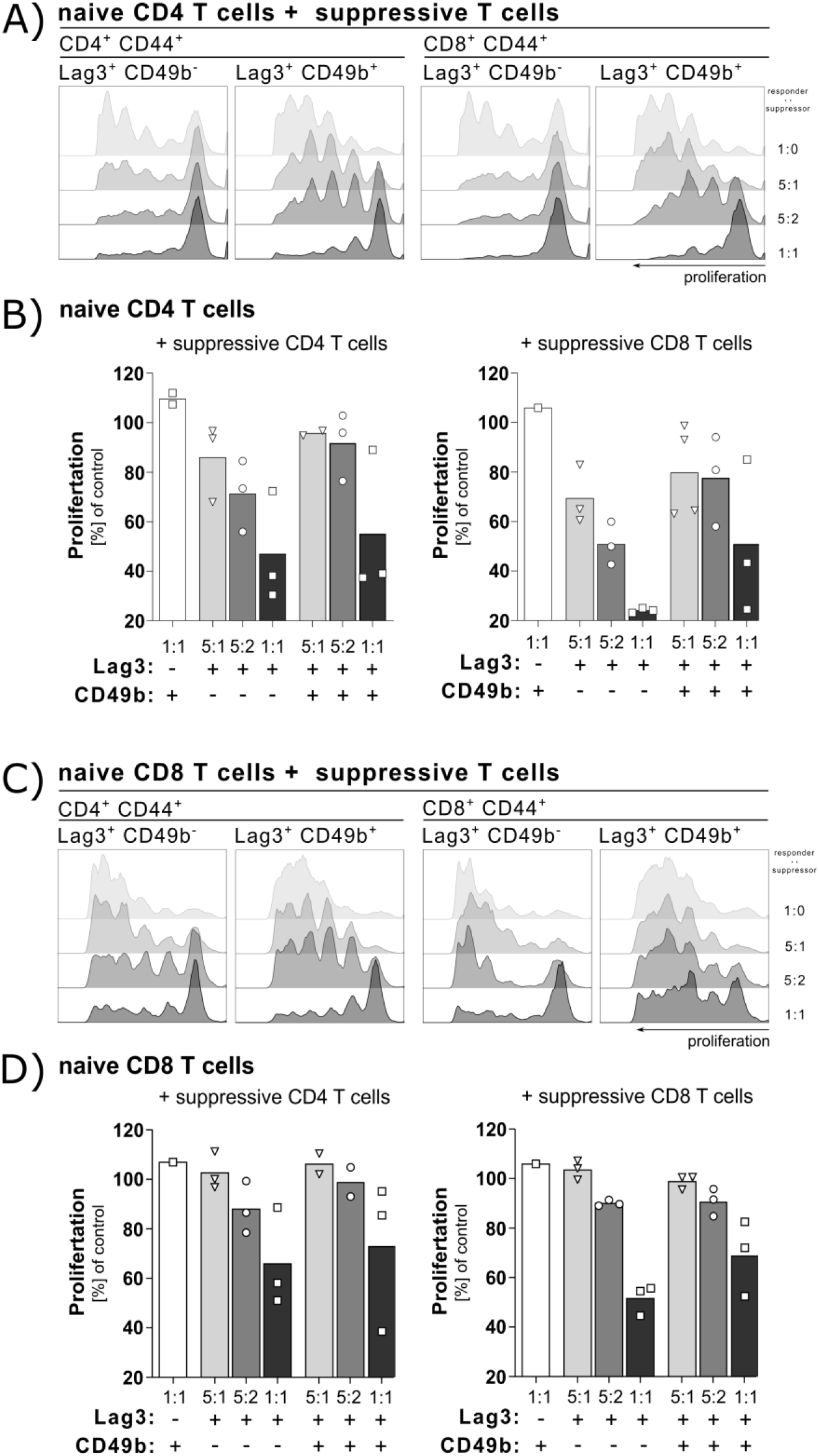
Malaria-induced T cells show regulatory capacity. PbA-induced potentially regulatory T cells were isolated 6 dpi and sorted for indicated subsets. Co-culture of effector T cells and potential regulatory T cells was performed for 3 days. Effector T cells were stained with eFluor450 to track proliferation. **(A)** Histogram of proliferated CD4^+^ T cells in indicated co-culture. Each panel shows a respective histogram out of 4-6 technical replications from 2-3 individual experiments. **(B)** Quantification of suppression of CD4^+^ effector T cells. Each dot represents one individual experiment with 2-3 technical duplicates. **(C)** Histogram of proliferated CD8^+^ T cells in indicated co-culture. Each panel shows a respective histogram from 2-3 technical repetitions out of 2-3 individual experiments. **(D)** Quantification of suppression of CD8^+^ effector T cells. Each dot represents one individual experiment with 2-3 technical repetitions.

As the induction of cytotoxic CD8^+^ T cells drives the immunopathology in malaria, we assessed whether PbA-induced regulatory LAG-3^+^ T cells could also suppress CD8^+^ T effector cells. *In vitro* suppression assays showed that malaria-induced CD4^+^LAG-3^+^ and CD8^+^LAG-3^+^ T cells could suppress CD8^+^ T cell proliferation. Again, the additional expression of CD49b did not lead to a more substantial suppressive capacity of LAG-3^+^ T cells.

In conclusion, malaria-induced LAG-3^+^CD4^+^ T cells show a suppressive effect on CD4^+^ and CD8^+^ T cells. Remarkably, we demonstrated that malaria-induced LAG-3^+^CD8^+^ T cells also displayed a suppressive capacity towards naïve CD4^+^ and CD8^+^ T cells.

### 4. Malaria-induced CD8^+^LAG-3^+^ T cells are still cytotoxic

The infection by PbA induces T cells co-expressing a variety of co-inhibitory molecules. Interestingly, we saw that the induced CD4^+^ and CD8^+^ T cells are not exhausted, but develop suppressive capacity. Further, we questioned whether induced CD8^+^ T cells are still cytotoxic, regardless of the expression of multiple inhibitory surface receptors and their suppressive capacity (Fig 6). For the implementation of an *in vitro* cytotoxic assay, we infected OT-I mice with a transgenic PbA (PbOVA) strain, expressing the MHCI epitope SIINFEKL (Lundie et al., 2008). All CD8^+^ T cells in the OT-I mouse strain are specific for SIINFEKL (Hogquist et al., 1994). We isolated the previously examined CD49b/LAG-3 CD8^+^ T cell subsets by cell sorting and co-cultivated PbOVA-induced sorted splenocytes with SIINFEKL-pulsed splenocytes. Surprisingly, the co-inhibitor-rich cells (here sorted as LAG-3 positive) demonstrated a higher cytotoxic capacity than their LAG-3^−^ counterparts by killing more antigen-loaded target cells in a dose-dependent manner (Fig 6B). Taken together, by investigating the expression of granzyme B, IFN_*γ*_, and IL-10, we demonstrated that malaria-induced CD4^+^LAG-3^+^ and CD8^+^LAG-3^+^ T cells are not exhausted, but rather highly activated. Moreover, malaria-induced T cells seem to develop a dual phenotype by simultaneously producing pro- and anti-inflammatory molecules.

**FIG 6:**
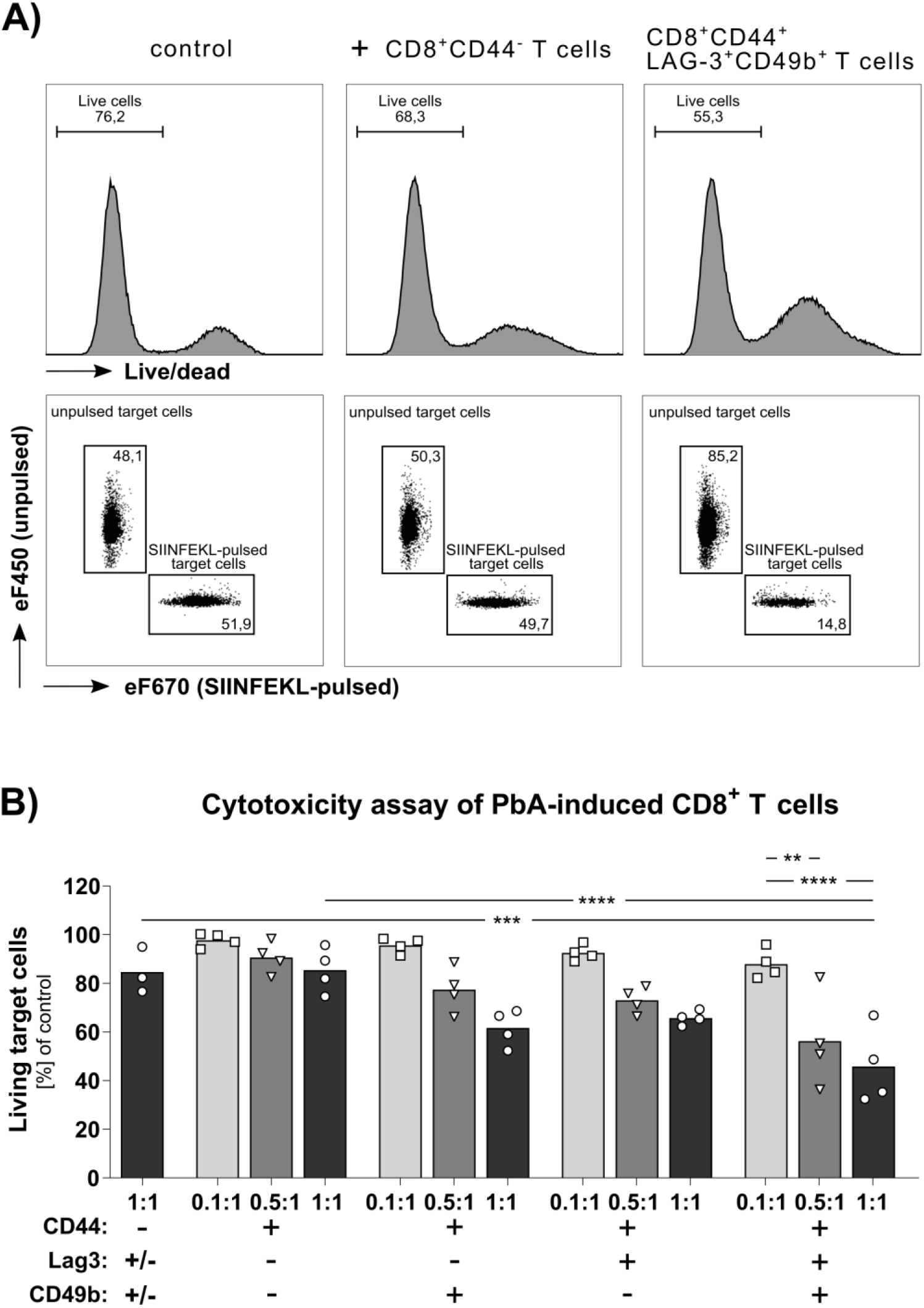
Malaria-induced regulatory CD8^+^ T cells are still cytotoxic. CD8^+^ T cells were isolated and sorted from PbOVA-infected OT-I mice 6 dpi and co-cultured with SIINFEKL-pulsed target cells to determine cytotoxic capacity of indicated subsets. **(A)** Part of the gating strategy to calculate the proportion of living SIINFEKL-loaded target cells after the co-culture with effector cells. Representative histograms and dot plots of a target cell control (target cells only, left), target cells co-incubated with CD8^+^CD44^−^ (middle), and CD8^+^CD44^+^CD49b^+^LAG-3^+^ T cells (right). **(B)** Antigen-specific killing of different CD8^+^ T cell subsets of OT-I mice infected with PbOVA. Ratios indicates ratio of effector to target cells in co-culture. Each dot represents an individual experiment with a minimum of 2 technical replicates. Ordinary one-way ANOVA with subsequent Tukey’s multiple comparison; P values ≤ 0.05 (*), ≤ 0.01 (**), ≤ 0.001 (***) or ≤ 0.0001 (****) were considered statistically significant.

### 5. CD4^+^LAG-3^+^ and CD8^+^LAG-3^+^ T cells are induced in patients with acute *P. falciparum* malaria

We showed that infection of C57BL/6 mice with PbA led to solid activation of adaptive immunity. A specific population of LAG-3^+^ T cells was induced, and these T cells were characterized by their expression of other co-inhibitory molecules like PD-1, CD39, TIGIT, or TIM-3. Interestingly, we observed that these cells also expressed granzyme B and IFN_*γ*_. Conversely, we demonstrated the expression of anti-inflammatory IL-10 by those malaria-induced T cells. Finally, we showed malaria-induced LAG-3^+^ T cells from the CD4 and CD8 subsets exert regulatory capacity towards CD4^+^ and CD8^+^ T cells. These results demonstrated the induction of potential regulatory CD4^+^ and CD8^+^ T cells in a mouse model of cerebral malaria.

To validate the expansion of these regulatory T cells in human malaria, we investigated blood samples of *P. falciparum*-positive travelers in Hamburg, Germany, treated at the University Medical Centre Hamburg Eppendorf. In concordance with the previous data in the murine PbA model, malaria patients showed a significantly higher frequency of CD4^+^LAG-3^+^ T cells than healthy donors (Fig 7A). Comparable to the induced regulatory T cells in the murine model, induced T cells in malaria patients expressed activation markers, like C-C chemokine receptor type 5 (CCR5) (Fig 7C+D), as well as surface markers associated with exhaustion, like PD-1, TIGIT, or TIM-3. Interestingly, CD4^+^LAG-3^+^ T cells expressed high amounts of the immune regulatory ectonucleotidase CD39 (Fig 7C).

**Fig. 7:**
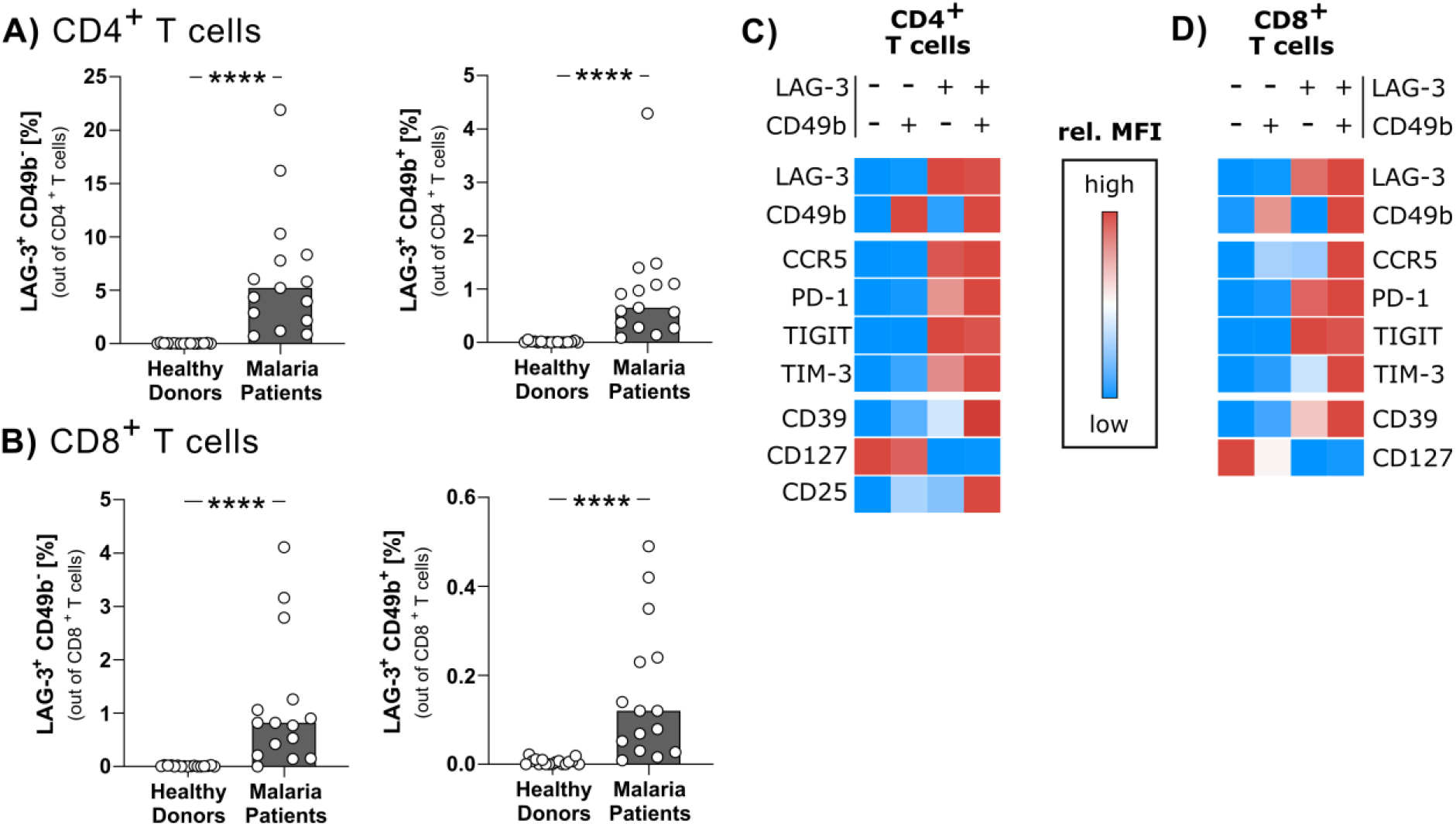
Malaria-induced LAG-3^+^ T cells in malaria patients. Whole blood samples of healthy donors and malaria patients were stained *ex vivo* and analyzed by flow cytometry. **(A)** and **(B)**: Frequencies of LAG-3^+^CD49b^−^ and LAG-3^+^CD49b^+^ cells within CD4^+^ and CD8^+^ T cells of healthy donors (n=17) and malaria patients (n=15). Bars represent the median. **(C)** and **(D)**: Relative MFI of surface markers of indicated LAG-3 and CD49b expressing CD4^+^ and CD8^+^ T cells of malaria patients (n=7-9). Mann-Whitney Test; p values ≤ 0.05 (*), ≤ 0.01 (**), ≤ 0.001 (***) or ≤ 0.0001 (****) were considered statistically significant.

Furthermore, when investigating CD8^+^ T cell subsets in malaria patients, we also observed the induction of CD8^+^LAG-3^+^ T cells (Fig 7B). Following our findings in the mouse model of PbA infection, induced CD8^+^LAG-3^+^ T cells were characterized by the expression of other co-inhibitory receptors like PD-1 and TIGIT (Fig 7C). In contrast to the mouse model, only the CD8^+^LAG-3^+^CD49b^+^ T cells also expressed TIM-3. Moreover, the CD8^+^LAG-3^+^CD49b^+^ T cells expressed higher levels of the activation marker CCR5 and the immune regulatory ectonucleotidase CD39 (Fig 7C+D) compared to their CD49b counterparts, indicating a higher suppressive capacity.

Collectively, by investigating the T cell response in malaria patients, these results show the induction of potentially regulatory CD4^+^ and CD8^+^ T cells in humans. Human LAG-3^+^CD49b^−^ and LAG-3^+^CD49b^+^ T cells share specific characteristics with the formerly described, PbA-induced, regulatory T cells in mice.

## Discussion

The blood stage of *P. falciparum* malaria is associated with robust activation of CD4^+^ as well as CD8^+^ T cells producing pro-inflammatory cytokines and cytotoxic molecules, respectively (Otterdal et al., 2018). Recent data suggest that an overwhelming T cell response contributes to a severe clinical outcome (Kaminski et al., 2019; Riggle et al., 2020), indicating that tight control of T cells is of significant importance for optimal resolution of a malaria episode. Thus, activation of T cells appears to be a tightrope walk between protection and pathology.

Acute malaria is associated with strong induction of co-inhibitory molecules on T cells. This was demonstrated in humans (Abel et al., 2018; Illingworth et al., 2013) as well as in animal models (Khandare et al., 2017; Lepenies et al., 2007). In chronic antigen exposure settings, expression of these molecules is associated with T cell exhaustion, especially of CD8^+^ T cells (Cornberg et al., 2013; Ochel et al., 2016). The function of these molecules in acute inflammatory settings is not well characterized, and it is not even clear whether these co-inhibitory molecules on CD4^+^ T cells are linked to their exhaustion as is described for CD8^+^ T cells. Using mouse models of acute malaria, it was already shown that blocking the exhaustion marker CTLA-4 could exacerbate the disease (Lepenies et al., 2007).

In contrast, a combined blockade of PD-1 and LAG-3 is associated with better control of parasitemia in a more chronic model of malaria (Butler et al., 2012). It is tempting to speculate that induction of co-inhibitory molecules in acute malaria regulates T cell activation. They either limit overwhelming T cell activation and immunopathology or limit parasite control in more chronic situations. Moreover, induction of co-inhibitory molecules might be, on the one hand, an immune escape mechanism of the parasite (Demarta-Gatsi et al., 2017) or, on the other hand, an emergency brake of the host preventing overactivation of the immune system (Cornberg et al., 2013). To further pinpoint the function of these molecules, a comprehensive picture of the induction and co-expression of these molecules on T cells and an association with their function is of crucial importance to further unravel immune regulatory networks in malaria.

To this end, we analyzed T cell activation after infection of C57BL/6 mice with *P. berghei* ANKA, a widely used model of experimental cerebral malaria, using the unbiased clustering approach FlowSOM (Van Gassen et al., 2015). These allowed us to compare T cells from different groups in an unbiased way, but on the same scaffold. Interestingly, a significant proportion of activated CD4^+^CD44^+^ as well as CD8^+^CD44^+^ T cells in the spleen and liver were PD-1^+^ and LAG-3^+^. Despite expressing these co-inhibitory molecules, they appeared to be fully functional T effector cells producing pro-inflammatory cytokines and cytotoxic molecules. Thus, CD8^+^CD44^+^LAG-3^+^ T cells were capable of lysing peptide-pulsed target cells.

Furthermore, LAG-3^+^ T cells were more cytolytic than their LAG-3^−^ counterparts. Thus, in acute malaria, the expression of co-inhibitory molecules delineates strongly activated T cells instead of being indicators of T cell dysfunction. However, we cannot exclude that the expression of co-inhibitory molecules restricts the function of these T cells or that in this acute model, with its very fast and robust induction of T cells, a prolonged expression would be needed to attenuate T cell exhaustion.

Interestingly, CD4^+^LAG-3^+^ and CD8^+^LAG-3^+^ T cells produce both pro-inflammatory cytokines like IFN_*γ*_ and anti-inflammatory cytokines like IL-10 and thus resemble Tr1 cells (Zeng et al., 2015). To further decipher their function, we analyzed their suppressive capacity and demonstrated that CD4^+^LAG-3^+^ T cells could suppress CD4^+^ and CD8^+^ T cells, regardless of CD49b expression. We also found that CD8^+^LAG-3^+^ T cells suppress CD4^+^ and CD8^+^ T cells, independent of CD49b expression. This is, to our knowledge, the first description of malaria-induced CD8^+^LAG-3^+^ T cells with suppressive capacity. Tr1 cells in chronic inflammatory conditions are characterized by co-expression of LAG-3 and CD49b (Gagliani et al., 2013). In our work, describing an acute disease, the suppressive capacities of LAG-3^+^CD49b^+^ and LAG-3^+^CD49b^−^ T cells are alike. However, as an integrin, CD49b might be associated with tissue localization and might not be functionally involved in their suppressive activity, and differences might be due to different disease models.

To confirm whether similar T cell subsets were induced in acute human malaria, we analyzed LAG-3-expressing T cells in acute *P. falciparum* malaria. Indeed, we found strong induction of LAG-3 on CD4^+^ and CD8^+^ T cells, and similar to the mouse model, a minor fraction of them co-express CD49b.

However, LAG-3^+^CD49b^+^ and LAG-3^+^CD49b^−^ CD4^+^ and CD8^+^ T cells are very similar regarding co-expression of other co-inhibitory molecules and express PD-1, TIGIT, and TIM-3. Thus, experiments to characterize the suppressive capacity of the different subsets of malaria-induced T cells expressing co-inhibitory receptors should be done. It is tempting to speculate that these T cells can modify the course of the disease by having a dual function in expressing pro-inflammatory cytokines and at the same time being suppressive to other T cells. It could be interesting to study these T cell populations in patients with either severe or uncomplicated course of the disease. Recently, we described the induction of CD4^+^ T cells in acute *P. falciparum* malaria co-expressing PD-1 and CTLA-4 (Mackroth et al., 2016). Interestingly, these cells can suppress other CD4^+^ T cells as efficiently as FoxP3^+^ Treg.

In summary, we show the induction of regulatory CD4^+^ T cells and CD8^+^ T cells during infection of C57BL/6 mice with PbA. These malaria-induced T cells co-express LAG-3 and several other co-inhibitory molecules, are capable of suppressing T cell proliferation, and share characteristics with Tr1 cells. Functional assays revealed that the LAG-3^+^CD8^+^ T cell subset could lyse antigen-pulsed target cells and suppress the function of other T cells. In addition, we provide evidence that T cells with similar characteristics are also induced in acute *P. falciparum* malaria in humans. This novel immune regulatory pathway may have evolved to prevent and/or control prolonged inflammation in the setting of strong immune activation as they are found in acute malaria.

## Methods

### Animals

C57BL/6J and BALB/c mice were bred at the animal facility of the BNITM, Hamburg, Germany in individual ventilated cages (IVC) with food and water *ad libidum.* All animal experiments were performed in accordance with German law and registered with the “Behörde für Gesundheit und Verbraucherschutz (BGV)” under license 83/17; O012/2018; N005/2019.

### *Plasmodium berghei* ANKA (PbA) Infection

Infected red blood cells (iRBC) were stored with 15 % glycerol (Merck, Darmstadt, Germany), 2.1 % sorbitol (Sigma Aldrich, St. Louis, Missouri) in PBS in liquid nitrogen. To achieve consistent blood-stage infections and avoid bias by frozen parasites, pre-experimental mice were infected with stabilate. Infected blood was obtained from pre-experimental mice after 5-7 days, and C57BL/6J mice were injected with 1×10^5^ iRBC in 200 μl PBS intraperitoneally (i.p.). For the generation of sporozoites, BALB/c mice were treated with 1.25 mg phenylhydrazine to stimulate erythrocyte generation two days before infection with 70-120 μl infected blood from pre-experimental mice. On day 3-4 post-infection, mice were fed as blood meals for 30 minutes to *Anopheles stephensi* mosquitos. After the blood meal, mosquitos were held at 21 °C, 80 % humidity, and a 14 h light /10 h dark day-night cycle. After 18-21 days, sporozoites were isolated from the salivary glands of infected mosquitos. Sporozoites were irradiated by exposure to a cesium-137 source. Age-matched (7-15 weeks) C57BL/6J were injected with 1000-5000 PbA sporozoites. Indicated groups were treated with 70 μg/ml pyrimethamine in drinking water 5 days after initial infection (Friesen et al., 2011).

### Parasitemia

PbA infection rate was determined by counting infected erythrocytes. A thin blood smear from infected mice was stained with Wright Giemsa Stain. A minimum of 1000 erythrocytes in min. 3 fields of views were counted.

### T cell isolation

Mice were euthanized, and 300-500 μl blood was drawn from the *vena cava* and mixed with 20 μl Heparin-Natrium-25000 (Ratiopharm, Ulm, Germany). Following the blood draw, mice were directly perfused with PBS/ 2 % FCS/ 1 mM EDTA and liver digest medium (ThermoFisher Scientific/ Gibco, Waltham, Massachusetts) through the *vena cava*. Leukocytes were isolated from blood, brain, liver, and spleen Centrifugations were done for 5 minutes, 400 x g at 4 °C unless otherwise stated.

#### Blood

Whole blood was taken from the *vena cava* and mixed with heparin. Erythrocytes were lysed with two times 20 ml lysis buffer (150 mM NH_4_CL, 10 mM KHCO_3_, 0.1 mM EDTA, pH 7.2-7.4). Erythrocytes-free single cells were directly used for staining.

#### Spleen

Prepared spleens were disrupted through a 100 μm sieve with PBS/ 2 % FCS/ 1 mM EDTA. The resulting suspension was centrifuged. Erythrocytes were lysed using lysis buffer. Cells were washed with PBS/ 2 % FCS/ 1 mM EDTA and filtered through another 100 μm cell strainer.

#### Liver

Lymphocytes from liver tissue were obtained as described previously (Ochel et al., 2016). Briefly, livers were cut into small pieces and digested. Single cells were separated from debris by 40 % Percoll (SIGMA, St. Louis, Missouri) in PBS/ 2 % FCS/ 1mM EDTA, and centrifugation was done for 20 minutes at 600 g at room temperature (RT). Erythrocytes were lysed.

#### Brain

The brain was strained through a 100 μm sieve, washed with PBS/ 2 % FCS/ 1 mM EDTA. Pellet was resuspended in 40 % Percoll in PBS/ 2 % FCS/ 1 mM EDTA and centrifuged at 600 g, RT. Erythrocytes were lysed before staining.

### Restimulation of lymphocytes

Cells were restimulated to induce cytokine production. Briefly, 2×10^6^ cells were incubated for 5 hours with Brefeldin A (BioLegend, San Diego, California) and Monensin (BioLegend, San Diego, California) at 37 °C, 5 % CO_2_. For restimulation, cells were incubated either with 1 mg/ml Phorbol 12-Myristat 13-Acetate (PMA) and 0.25 mg/ml Ionomycin (Iono) or 1 μg/ml PbA-peptide mix (SQLLNAKYL; IITDFENL; EIYIFTNI from Jerini Biotools (Poh et al., 2014)) or left unstimulated.

### Suppression assay

To investigate the suppressive capacity of the different T cell subpopulations, suppression assays were performed. Splenocytes from naïve C57BL/6 mice were obtained as described previously. CD4^+^ or CD8^+^ T cells (target cells) were obtained using the respective negative selection kit (Miltenyi, Bergisch Gladbach, Germany) and were labeled with 1 μM of proliferation dye eFluor450 (ThermoFisher, Waltham, Massachusetts). Potential suppressive T cell populations were obtained from spleens of infected C57BL/6 mice at day 6 post-infection and sorted using a FACS Aria II (BD Bioscience, Franklin Lakes, New Jersey). 5×10^4^ naïve target CD4^+^ or CD8^+^ T cells were seeded in complete RPMI 1640 (RPMI 1640 + 5 % FCS + 2 mM L-glutamine + 10 mM HEPES + 50 μg/mL Gentamicin) in a round bottom 96 well plate and co-cultivated with different ratios of potential suppressor T cells (1:0, 5:1, 5:2 and 1:1 target; suppressor T cells). The cells were cultured with αCD3/αCD28 beads (Invitrogen, Carlsbad, California) at a ratio of 1:1 (beads: total T cells) and incubated at 37 °C, 5 % CO_2_ for 3 days.

### CD8^+^ cytotoxicity assay

CD8^+^ T cells were isolated and sorted as described above from spleens of OT-I mice infected with PbOVA. Splenocytes from naïve C57BL/6 mice were split into two groups, one of which was pulsed with 1 μg/ml SIINFEKL peptide in complete RPMI 1640 for 1 h at 37 °C, 5 % CO_2_. The groups (pulsed and unpulsed) were labeled with eFluor450 or eFluor670 (ThermoFisher, Waltham, Massachusetts), respectively and then mixed. Mixed target cells were co-incubated with the sorted CD8^+^ T cells in indicated ratios for 6 h. Afterwards, living cells were identified by ZombieNIR Live/Dead staining (BioLegend, San Diego, California). By calculating the proportion of SIINFEKL-loaded target cells from the remaining living target cells compared to a control without any sorted CD8^+^ T cells, the cytotoxic capacity of each sorted subset was determined.

### Flow cytometry analysis of murine cells

The following antibodies were used for surface staining at RT for 20 min: αCD3 BUV395 (145-2C11), αCD4 V500 (RM4-5), αCD8 V450 (53-6.7), and αCD223 (LAG-3) BUV737 (C9B7W) purchased from BD Bioscience, Franklin Lakes, New Jersey. αCD25 AlexaFluor 700 (PC61), αCD27 FITC (LG4.A10), αCD39 PE-Cy7 (Duha59), αCD44 AlexaFluor 700 or BV650 (IM7), αCD49b PE or AlexaFluor 488 (DX5), αCD62L PE-Cy5 or PerCP (MEL-14), αCD69 BV785 (H1.2F3), αCD223 (LAG-3) PerCP-Cy5.5 (C9B7W), αCD278 (C398.4A), αCD279 (PD-1) APC or PE-Cy7 (RMP1-39), αCD366 (Tim-3) BV711(RMT3-23) and αTIGIT PE-Dazzle594 (2G9) were purchased from BioLegend, San Diego, California and αKLRG-1 APC-eFluor780 (2F2) was purchased from ThermoFisher, Waltham, Massachusetts. According to the manufacturer’s instruction, intracellular staining was performed using the FoxP3 transcription factor staining kit (eBiosience, San Diego, California). Fixation/ permeabilization was followed by intracellular staining for 20 minutes at RT. αgranzyme B Pacific blue (GB11, BioLegend, San Diego, California), αCD3 BUV395, and αFoxP3 PerCP-Cy5.5 (FJK-16s, ThermoFisher, Waltham, Massachusetts) were used. Samples were acquired on a BD LSRII or BD LSRFortessa (BD Bioscience, Franklin Lakes, New Jersey). FACS data were analyzed using FlowJo v10 (BD Bioscience, Franklin Lakes, New Jersey).

### Data processing and clustering

Lymphocytes were cleaned from doublets and gated for CD3 to identify T cells. CD3^+^ T cells were subsequently gated for the expression of CD4 and CD8. CD8^+^ T cells or CD4^+^ T cells were downsampled, and respective groups were concatenated (2-3 individual mice per group). Clustering was performed by FlowSOM (package v2.6; random starting seeds) in FlowJo (Van Gassen et al., 2015). In total, 15 markers were included (LAG-3, TIM-3, TIGIT, PD-1, CD39, FoxP3, CD25, CD27, CD49b, granzyme B, KLRG-1, ICOS, CD69, CD62L, CD44). The grid size of the self-organizing map (SOM) was set to 15×15. Metacluster number was set to 15. Concatenated expression data from splenocytes from PbA-infected animals and naïve controls were used to generate a master cluster grid and a minimum-spanning tree (MST). To identify differences in the expression pattern, the individual groups were computed based on the master grid. CD4 and CD8 T cells were analyzed individually.

### Human study population

In total, 15 malaria patients and 17 healthy adults between the age of 18-61 were enrolled in this study. All malaria patients were returning travelers infected with *P. falciparum* in a malaria-endemic area, but developed their malaria symptoms after returning to Germany. The malaria patients were enrolled during their malaria treatment at the University Medical Centre Hamburg Eppendorf (Hamburg, Germany). The healthy adults were enrolled at the Bernhard Nocht Institute for Tropical Medicine in Hamburg. Ethical approval was obtained for the study from the Ethics Committee Hamburg, Germany (PV 5537). Written informed consent was given from all study participants before inclusion in the study.

**Tab. 1.** Clinical data of enrolled malaria patients.

| # | Sex | Age | Country<br>of<br>infection | Parasitemia<br>at the day of<br>diagnosis | Parasitemia at the<br>day of sampling | Day (sampling)<br>after the start<br>of treatment |
| --- | --- | --- | --- | --- | --- | --- |
| 1 | f | 47 | Togo | 5 % | negative | 2 |
| 2 | m | 45 | Togo | < 1 % | < 1 % | 3 |
| 3 | f | 26 | Togo | < 1 % | negative | 3 |
| 4 | m | 26 | Guinea-Bissau | < 1 % | < 1 % | 3 |
| 5 | m | 41 | Nigeria | < 1 % | negative | 3 |
| 6 | m | 33 | Benin | < 1 % | negative | 4 |
| 7 | m | 43 | n.a. | 2.50 % | negative | 2 |
| 8 | f | 18 | Cameroon | 1.50 % | < 1 % | 2 |
| 9 | f | 61 | Tanzania | 21 % | negative | 16 |
| 10 | m | 49 | Nigeria | < 1 % | < 1 % | 1 |
| 11 | f | 39 | Cameroon | 6% | negative | 4 |
| 12 | m | 51 | Ivory<br>Coast | 0.50 % | negative | 4 |
| 13 | m | 50 | Togo | 0.50 % | < 1 % | 3 |
| 14 | m | 41 | Togo | < 1 % | negative | 4 |
| 15 | f | 55 | Ghana | 5% | negative | 3 |
|  | f/m =<br>6/9 | mean=<br>41.7 |  |  |  |  |

**Tab. 2.** Clinical Data of healthy donors.

| # | Sex | Age |
| --- | --- | --- |
| 1 | f | 59 |
| 2 | f | 27 |
| 3 | m | 27 |
| 4 | m | 26 |
| 5 | m | 34 |
| 6 | f | 25 |
| 7 | f | 57 |
| 8 | f | 51 |
| 9 | f | 27 |
| 10 | m | 35 |
| 11 | f | 31 |
| 12 | m | 29 |
| 13 | m | 53 |
| 14 | f | 26 |
| 15 | f | 24 |
| 16 | f | 21 |
| 17 | f | 39 |
|  | f/m= 11/6 | mean = 34.8 |

### Determination of *Plasmodium falciparum* infection

*P. falciparum* infection and parasitemia were determined by thick and thin blood film stained with Giemsa and examined with an oil immersion microscope (100x magnification).

### *Ex vivo* staining of human peripheral blood

Venous blood was collected into Lithium-Heparin tubes and processed within 5 hours. 100 μl of whole blood were incubated at 4 °C for 30 minutes with the following surface antibodies: CD3 APC/Cy7 (HIT3a), CD4 BV510 (RPA-T4), CD8 AF700 (RPA-T8), CD127 AF488 (A019D5), CD25 PE/Cy7 (BC96), PD-1 PerCP/Cy5.5 (EH12.2H7), TIGIT BV605 (A15153G), TIM-3 BV650 (F38–2E2), CD39 PE/Dazzle594 (A1), CD49b FITC (P1E6-C5), CCR5 PE (J418F1) (all BioLegend, San Diego, California) and LAG-3 APC (3DS223H) (eBioscience, San Diego, California). Afterwards, the cells were lysed and fixed with 2 ml of 1x RBC Lysis/Fixation Solution (Biolegend) for 15 minutes at RT and washed with cold FACS-buffer (PBS with 1 % FCS). Cells were analyzed on a LSRFortessa (BD Bioscience, Franklin Lakes, New Jersey) or a CytoFLEX S (Beckman Coulter, Brea, California) flow cytometer and analyzed with FlowJo v10 (BD Bioscience, Franklin Lakes, New Jersey). Lymphocyte doublets were excluded by FSC-A/FSC-H gating. Afterwards, T cells were gated by their expression of CD3. CD3^+^ T cells were subsequently gated on CD4 and CD8. CD4^+^ and CD8^+^ T cells were gated for LAG-3 and CD49b. LAG-3^+/−^CD49b^+/−^ populations were gated to determine the median fluorescence intensities (MFI) of all analyzed surface markers on these populations. This was done separately for CD4^+^ and CD8^+^ T cells, respectively. According to fluorescence minus one (FMO) controls, gates were set to identify false positives and spillover.

### Statistics

Mouse data were checked for normality by the D’Agostino & Pearson test or the Shapiro-Wilk test for small group size. If a Gaussian distribution is assumed, ordinary one-way ANOVA with subsequent Tukey’s multiple comparison test was performed. Otherwise, the Kruskal-Wallis test with Dunn’s multiple comparisons test was performed. Comparison of just two groups was made by Mann-Whitney test. Statistic calculations were done with Graphpad Prism v8 (Graphpad Software, San Diego, USA).

For human samples, frequencies of different T cell subsets of the two groups were compared using unpaired, nonparametric Mann-Whitney tests (GraphPad Prism v8).

P values ≤ 0.05 (*), ≤ 0.01 (**), ≤ 0.001 (***) or ≤ 0.0001 (****) were considered statistically significant.

## Financial support and competing interests

### The authors declare no competing interests

This work was funded via the Collaborative Research Center 841 of the German Research Foundation (SFB 841), 53170 Bonn, Germany. MM was funded by a DZIF clinical leave stipend (German Centre for Infection Research).

